# Identification and localization of bone marrow ILC2s using Id2/Cxcr6 dual-reporter mice

**DOI:** 10.64898/2026.11.09.765290

**Authors:** Cassandra Lepetit, Paloma Hedde, Godefroy Jacquemin, Elisa Gomez Perdiguero, Stéphane J.C. Mancini, Rachel Golub

## Abstract

Type 2 innate lymphoid cells (ILC2s) are present in the bone marrow, where their identification and spatial localization remain challenging because of their low abundance and the expression of shared markers by other lymphoid populations. Here, we evaluated the use of Id2^YFP/+^ Cxcr6^GFP/+^ dual-reporter mice to identify and visualize bone marrow ILC2s. Using spectral flow cytometry, we showed that both ST2⁻ and ST2⁺ ILC2s expressed high levels of ID2, whereas CXCR6 expression differed between these subsets. However, the ID2 expressing compartments also contained NK cells, NKT cells, conventional T cells and ILC progenitors. Surface marker profiling identified CD25 and ICOS as useful markers for enriching ILC2 populations, with ST2 enabling the segregation of the ST2^-^ and ST2⁺ subsets. Single-cell RNA sequencing further characterized the diversity of Id2 and Cxcr6 expressing lymphoid populations and supported the use of CD5 and NKp46 as exclusion markers. Combining these markers with the dual-reporter system, confocal microscopy revealed YFP⁺GFP⁺CD5^-^NKp46^⁻^ cells compatible with an ILC2 phenotype, distributed throughout the bone marrow as isolated cells or within multicellular clusters. Complementary flow cytometric analyses demonstrated that ILC2s accounted for more than 90% of a phenotypically related reporter-positive population after exclusion of major T and NK cell populations, providing additional support for the identification of the cells visualized in situ as putative ILC2s. Together, these findings establish a strategy combining dual-reporter mice, flow cytometry, transcriptomic profiling and confocal imaging to investigate the localization and spatial organization of putative bone marrow ILC2s.

**Highlights:** • Bone marrow ILC2 subsets differ in CXCR6 expression but uniformly express high ID2 levels.

• ID2 and CXCR6 are co-expressed by several bone marrow lymphoid populations.

• CD25, ICOS and ST2 improve the identification of bone marrow ILC2 subsets.

• Single-cell RNA-seq supports CD5 and NKp46 as exclusion markers.

• Dual-reporter mice combined with exclusion markers enable imaging of putative BM ILC2s, supported by flow cytometric validation.

## Introduction

Type 2 innate lymphoid cells (ILC2s) are tissue-resident cells distributed across several organs, including adipose tissue, the lung, the intestine and the bone marrow (BM)[1]. They play crucial roles in orchestrating type 2 immune responses, contributing to allergic inflammation, parasite clearance, and tissue repair [2–4]. ILC2s arise from the common lymphoid progenitor (CLP), which resides in the fetal liver and BM and gives rise to all lymphoid lineages [5–8]. ILC development involves progenitor populations including the early innate lymphoid progenitor (EILP) stage and subsequently the common helper-like innate lymphoid progenitor (ChILP) [9–11]. Downstream of the ChILP stage, innate lymphoid cell precursors (ILCPs), retain the capacity to give rise to ILC1, ILC2, and ILC3 subsets [12]. A defining feature of ILC lineage commitment is the requirement for the transcriptional repressor ID2, an inhibitor of E-protein transcription factors [13–16]. Although ID2 is essential for ILC development, its expression is not restricted to the ILC lineage, limiting its use as a specific marker for identifying ILCs within the heterogeneous BM environment [17,18].

Recent studies using sophisticated polychromatic reporter mice have provided unprecedented resolution of ILC precursor heterogeneity, firmly establishing that commitment to the ILC2 lineage is marked by the co-expression of the transcription factors Bcl11b and Gata-3 [19,20]. While ID2 is essential for early ILC development, its role in adaptive lymphocytes reveals complex, lineage-specific functions. In CD8^+^ T cells, ID2 is upregulated during immune responses where it is critically required for the survival of effector cells, thereby enabling the development of a robust memory pool [21]. This role in enforcing a differentiated state extends to CD4⁺ T cells, where high ID2 expression drives pathogenic effector functions in Th1 and Th17 responses [22,23] and regulates the differentiation program of tissue-resident memory T cells [24]. Thus, ID2 expression in multiple lymphoid populations represents a major challenge for the specific identification of ILC2s using reporter-based approaches.

Previous studies have demonstrated that tissue-resident progenitors contribute to the establishment and maintenance of peripheral ILC populations, while the BM remains an important site of ILC development and differentiation [1,25]. Notably, ILC2s represent a prominent mature helper-like ILC population in the BM, whereas ILC1 and ILC3 populations are poorly represented or difficult to identify under steady-state conditions [25]. These observations suggest that BM ILC2s constitute a specialized resident population potentially involved in the regulation of the hematopoietic microenvironment. BM ILC2s have been reported to regulate hematopoietic stem cell proliferation and myeloid differentiation through IL-33/ST2-dependent production of IL-6 and GM-CSF, particularly in the context of hematopoietic stress and aging [26] and to promote hematopoietic recovery after 5-fluorouracil treatment via GM-CSF production [27].

Although BM ILC2s contribute to hematopoietic regulation, their localization and spatial organization within the marrow remain poorly characterized. To address this gap, we characterized ID2 and CXCR6 reporter expression among BM lymphoid populations to develop a strategy for visualizing putative BM ILC2s in situ using ID2^YFP/+^ CXCR6^GFP/+^ dual-reporter mice. This dual-fluorescent system simultaneously reports ID2 and CXCR6 expression through YFP and GFP fluorescence, respectively [28]. CXCR6 has been identified as a marker highly expressed by ILC precursor populations in the fetal liver [29] as well as by ILC progenitors in the adult BM [28]. However, both ID2 and CXCR6 are expressed by other lymphoid populations, their combined expression alone being thus insufficient to specifically identify BM ILC2s.

We first characterized the distribution of ID2 and CXCR6 reporter expression among BM ILC2s and other lymphoid populations using spectral flow cytometry. We then combined phenotypic and transcriptomic analyses to identify additional markers capable of distinguishing ILC2s from other reporter-positive cells. Finally, we applied this strategy to confocal imaging of BM sections, enabling the visualization of ILC2 compatible cells within their native tissue environment and the characterization of their spatial distribution.

## Materials and Methods

### Mice

ID2^YFP/+^ CXCR6^GFP/+^ mice were in a full C57BL/6J background and were maintained under specific-pathogen-free conditions, air renewal (20 times per hour), constant ambient temperature of 22 ± 2C and 14:10 h light-dark cycle. All animal procedures were approved by the Pasteur Institute Safety Committee and conducted according to French and European Community institutional guidelines.

### Cell preparation

BM was isolated by flushing the hindlimbs with Hanks’ balanced salt solution (HBSS) (Gibco) supplemented with 2% FCS (Gibco). Red blood cells were lysed (1 min) with Ammonium-Chlorure-Potassium (ACK) solution (Gibco).

### Flow cytometry and cell sorting

For lineage exclusion, BM cells were first stained with a cocktail of biotin-conjugated antibodies against CD19 (6D5), Ter119 (Ter119), Gr-1 (RB6-8C5), and CD11b (M1/70), followed by incubation with streptavidin-PE-Cy5. Lineage-positive cells were excluded from all analyses. After exclusion staining, cells were incubated with directly conjugated antibodies specific for the following mouse antigens and fluorochromes: fluorescein isothiocyanate (FITC), phycoerythrin (PE), PECy5, PE-CF594, PECy7, allophycocyanin (APC), Alexa Fluor 700, APC-Fire810, APC-eFluor 780, Spark Red 718, BV421, BV510, BV570, BV605, BV650, BV711, BV786, BUV395, BUV496, BUV563 and BUV805 and were specific for the following mouse antigens: CD3e (145-2C11), CD19 (6D5), NK1.1 (PK136), CD8 (53-6.7), CD4 (RM4-5), Thy1.2 (30-H12), NKp46 (29A1.4), CD45 (30-F11), ST2/IL33R (DIH9), CD127/IL-7Rα (A7R34), Sca-1 (D7), ICOS (C398.4A), IL-17RB/IL-25R (MUNC33), LPAM1/α4β7 (DATK32), CD25 (PC61). Flow cytometry data were acquired using a Cytek Aurora, unmixed with SpectroFlo and analysed with Flow Jo sowftwares.

## Single cell RNA sequencing

Mice were euthanized, femurs and tibias were collected and flushed with FACS buffer (PBS, 1% Fetal Calf Serum, 2mM EDTA). Red blood cells were lyzed with ACK buffer. The lymphoid cells were stained with B220 (clone: RA3-6B2; Cat: 612839; BDBiosciences), CD3 (clone: 145-2C11; Cat: 100349; Biolegend), NK1.1 (clone: PK136; Cat: 108704; Biolegend), CD138 (clone: 281-2; Cat: 142505; Biolegend), CD2 (clone: RM2-5; Cat: 100114; Biolegend) and myeloid cells excluded using CD11c (clone: N418; Cat: 117372; Biolegend) and Gr1 (clone: RB6-8C5; Cat: 557979; BDBiosciences) before sorting by flow cytometry using a FACS AriaFusion (BD Biosciences).

Lin^-^Sca1^+^CD117^+^ (LSK) cells, CD127^+^ lymphoid progenitors and ILC2s were stained following depletion of Lineage positive cells with the direct lineage depletion kit (Miltenyi).

Cells were stained with CD117 (clone: 2B8; Cat: 105847; Biolegend), Sca1 (clone: D7; Cat: 741708; BDBiosciences), Ly6D (clone: 49-H4; Cat: 561147; BDBiosciences), CD127 (clone: A7R34; Cat: 135006; Biolegend), ST2 (clone: U29.93; Cat: 566309; BDBiosciences), CD90 (clone: 53-2.1; Cat: 565257; BDBiosciences), Lineage markers (Ter119 (clone: TER119; Cat: 116223; Biolegend), B220 (clone: RA3-6B2; Cat: 47-0452-82; ThermoFisher), CD19 (clone: 6D5; Cat: 115578; Biolegend), CD11b (clone: M1/70; Cat: 557657; BDBiosciences), CD71 (clone: R17217; Cat: 47-0711-82; ThermoFisher)) and sorted by flow cytometry.

Dead cells were stained with Sytox Blue (Thermo) and gated out during the sortings.

The libraries were generated using the Chromium Next GEM Single Cell 5’ Reagent Kits v2 (10x Genomics). Briefly, lymphoid cells on one side and LSK/lymphoid progenitors/ILC2s on the other were resuspended in the master mix and loaded together with partitioning oil and gel beads into the chip to generate the gel-bead-in-emulsion (GEM). The poly-A ARN from the cell lysate contained in every single GEM was retrotranscripted to cDNA, which contains an Illumina R1 primer sequence, Unique Molecular Identifier (UMI) and the 10x Barcode. The pooled barcoded cDNA was then cleaned up with Silane DynaBeads, amplified by PCR and the appropriately sized fragments were selected with SPRIselect reagent for subsequent library construction. During the library construction Illumina R2 primer sequence, paired-end constructs with P5 and P7 sequences and a sample index were added. Libraries were sequenced on the NovaSeqX platform.

## Single cell RNA-seq analysis

The demultiplexing, barcoded processing, gene counting and aggregation were made using the CellRanger v8.1.1 software. Cells were filtered based on the following parameters for both datasets: feature counts between 900 and 7500; reads related to mitochondrial genes below 10%; ribosomal reads above 5%. After an initial UMAP clustering, clusters corresponding to T cells, NK cells, NKT cells were defined using specific markers and selected for further analysis (total cell number = 2873). The cluster corresponding to ILC2s (235 cells) was similarly extracted from the “lineage negative cells” dataset and merged to the other cleaned dataset for a total of 3108 cells. The datasets were normalized using the harmony R package before the UMAP clustering.

The different lymphoid subsets were defined as follow: CD8 effector memory T cells (Ccl5^+^, Cxcr3^+^, Ccr7^lo^); CD8 central memory T cells (Lef1^+^, Tcf7^+^, Ccr7^+^, Sell^+^); CD8 resident memory T cells (Ccl5^+^, Cxcr6^+^, Pdcd1^+^); CD8 virtual memory T cells (Ly6a^+^, Nkg7^+^, Il2rb^+^, Klrk1^+^, Eomes^+^); CD8 stem-like memory T cells (Lef1^+^, Tcf7^+^, Bcl2^+^, Sell^+^, Slamf6^+^); CD4 naive T cells (Tcf7^+^, Lef1^+^, Ccr7^+^, Sell^+^, Cd44^-^); CD4 regulatory T cells (Foxp3^+^, Ctla4^+^, Nrp1^+^); double negative T cells (Cd4^-^, Cd8a^-^, Cd8b1^-^); Vγ1^+^ γδT cells (Trgv1^+^); γδTh17 (Trgv5^+^, Rorc^+^, Il23r^+^, Ccr6^+^); Immature NKT (Cd3e^+^, Sell^+^, Cd69^lo^, Klrbc^lo^); NKT1 (Cd3e^+^, Nkg7^+^, Klrb1c^+^, Ifng^+^, Tbx21^+^); NKT2 (Cd3e^+^, Zbtb16^+^, Cxcr6^+^); Immature NK (Ncr1^+^, Il2rb^+^, Id2^+^, Itgam^-^, Zeb2^-^); mature NK (Ncr1^+^, Id2^+^, Klra4^+^, Prf1^+^, Gzmb^+^, Eomes^+^, Zeb2^+^, Itgam^+^); ILC2 (Gata3^+^, Icos^+^, Il1rl1^+^, Il17rb^+^).

To improve the resolution of the different CD4 T cell subsets, a final sub-clustering was performed on the following clusters: Tregs, naive T cells, CD8 Trm, CD8 Tcm and Vγ1^+^ γδT cells.

## Immunofluorescence imaging of femur

Femurs were fixed in Antigenfix (Diapath France) 24h at 4°C. For cryopreservation, the bones were incubated sequentially in 10%, 20%, and 30% sucrose/PBS at 4°C for 1 hour each and embedded frozen in SCEM embedding medium (Section-Lab). Frozen sections were prepared at 20 μm thickness with a cryostat using the Kawamoto’s tape transfer method [30]. For immunofluorescence staining, sections were post-fixed with cold Antigenfix for 10 min and rinsed with PBS, followed by blocking with 20% donkey serum (DS; Biowest) in 0.1% Triton X-100/PBS for 1 hour at room temperature. Primary antibodies for CD5 (clone: 53-7.3; Cat: 100634; Biolegend), NKp46 (goat polyclonal antibody; Cat: AF2225; Biotechne), CD90 (clone: G7; Cat: 14-0901-82; Invitrogen), Endomucin (clone: V.7C7; Cat: sc-65495; Santa Cruz) and Sca-1 (clone: D7; Cat: 108112; Biolegend) were diluted in 2% DS 0.01% Triton X-100/PBS and incubated overnight at 4°C in a humid chamber. Primary antibody staining was followed by three washes with PBS and a 1-hour incubation with Alexa Fluor 594 and/or Alexa Fluor 647-conjugated secondary antibodies (Jackson ImmunoResearch). Slides were then incubated with 10 µM DAPI (Abcam) for nuclei staining. Slides were mounted in mowiol and dried overnight at room temperature before acquisition.

For whole-mount femur imaging, the BM was exposed by shaving the bone using a cryostat. Shaved femurs were fixed 30 min at 4°C in Antigenfix. Samples were then incubated in saturation buffer (0.1% Triton X-100/PBS-DS 20%) during 1 hour at room temperature.

Primary antibodies were incubated 72 hours at 4°C in 2% DS 0.01% Triton X-100/PBS. Secondary antibodies and DAPI were incubated 24 hours at 4°C in PBS.

All images were acquired using a confocal SP8 microscope (Leica) and visualized with Napari viewer for 3D volumes (https://napari.org/stable/index.html) or Fiji (ImageJ) for z-stack projections.

The number of CD5^+^, NKp46^+^ and CD5^-^NKp46^⁻^ cells among the Id2-YFP^+^Cxcr6-GFP^+^ double-positive (DP) cells was counted manually across z-stack images using QuPath software. (https://qupath.github.io). Images of femoral BM from three different mice were analyzed for a total count of 99, 79 and 100 DP cells respectively. The clusters were defined as an aggregate composed of at least 10 YFP^+^, GFP^+^ and/or YFP^+^GFP^+^ cells with a minimum of 5 YFP^+^GFP^+^ cells per cluster. The smallest cluster measured approximately 45 µm of diameter. A minimum of two clusters per bone marrow sections were observed.

## Results

### Identification of BM ILC2s within heterogeneous ID2 and CXCR6 expressing lymphoid populations

Previous studies using reporter mouse models have established that ID2 and CXCR6 can be used to characterized ILCs and their progenitors [10,19,20]. To develop a strategy for visualizing and identifying BM ILC2s by confocal microscopy using Id2^YFP/+^ Cxcr6^GFP/+^ dual-reporter mice, we sought to identify surface markers that could distinguish ILC2s from other lymphoid populations sharing these reporter profiles. We therefore used spectral flow cytometry to characterize the cellular composition of the ID2^hi^ CXCR6⁻ and ID2^hi^ CXCR6^hi^ compartments and identify candidate surface markers for distinguishing ILC2s in BM tissue sections (**Figure 1**).

**Figure 1.**
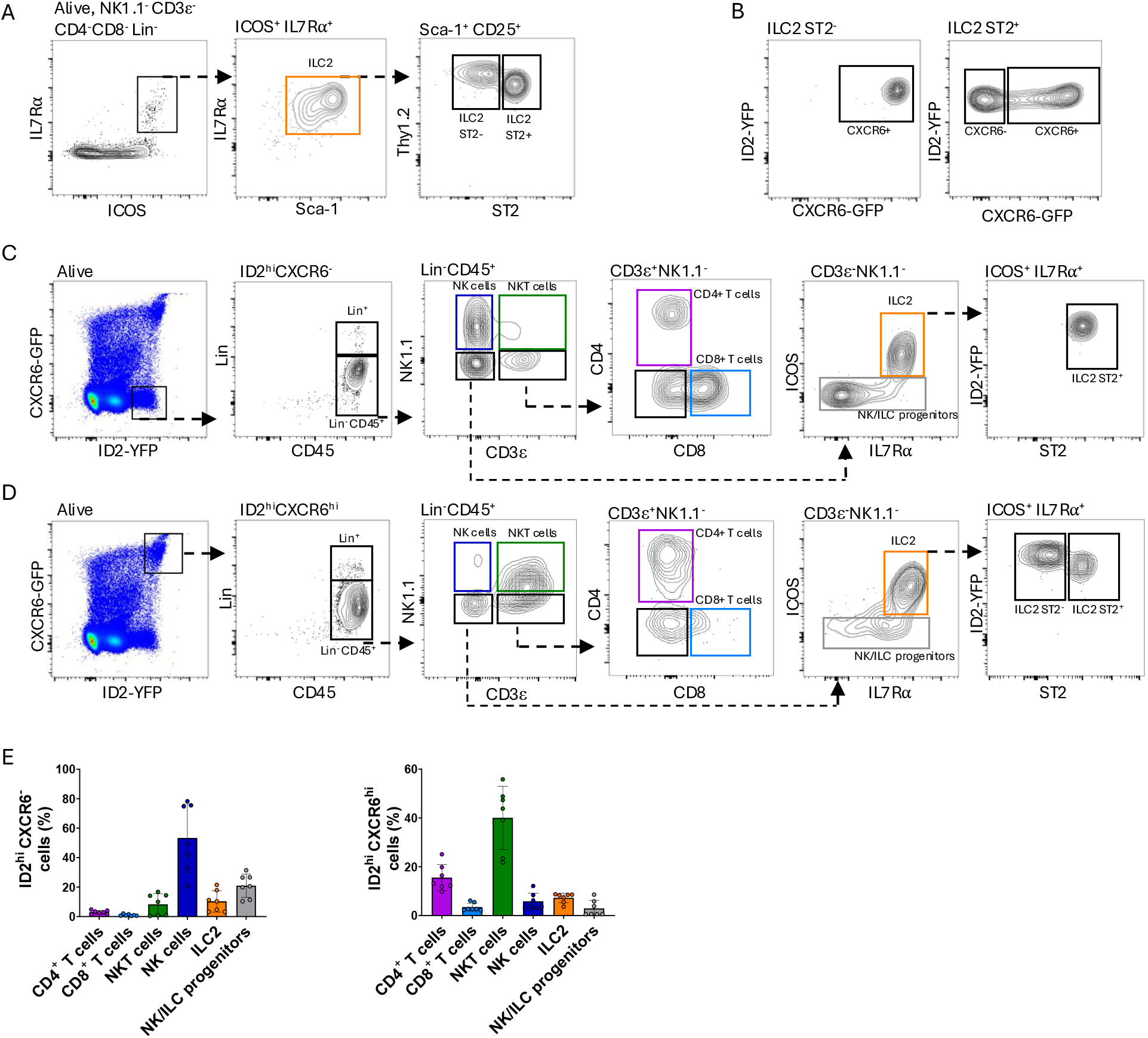
Identification of BM ILC2 subsets within heterogeneous ID2 and CXCR6expressing lymphoid populations. (A) Gating strategy for the identification of BM ST2⁻ and ST2^+^ ILC2 subsets among live NK1.1⁻CD3ε⁻CD4⁻CD8α⁻Lin⁻ (CD11b, Gr-1, CD19, Ter119) cells. (B) Representative flow cytometry plots showing ID2-YFP and CXCR6-GFP expression in gated ST2⁻ and ST2^+^ ILC2 populations. (C, D) Representative flow cytometry plots showing the distribution of immune cell populations within (C) ID2ʰⁱCXCR6^-^ and (D) ID2ʰⁱCXCR6ʰⁱ gates. (E) Quantification of the indicated populations within the total Lin⁻ID2ʰⁱCXCR6^-^ and Lin⁻ID2ʰⁱCXCR6ʰⁱ compartments. Data are presented as mean ± SD. Data were obtained from two independent experiments involving a total of seven mice.

BM ILC2s were identified as Lin⁻ (CD11b⁻, Gr-1⁻, CD19⁻, Ter119⁻), CD3ε⁻, CD4⁻, CD8α⁻, NK1.1⁻, ICOS⁺, IL-7Rα⁺, Thy1.2⁺ and Sca-1⁺ cells and further subdivided according to ST2 expression (**Figure 1A**). Both ST2⁻ and ST2⁺ ILC2 subsets exhibited high ID2 reporter expression. However, their CXCR6 reporter expression differed with ST2⁻ ILC2s being uniformly CXCR6^hi^, whereas ST2⁺ ILC2s comprising both CXCR6⁻ and CXCR6⁺ populations, present at approximately equal frequencies (**Figure 1B**). Thus, the combination of ID2 and CXCR6 reporters enables the detection of all ST2⁻ ILC2s and a substantial fraction of ST2⁺ ILC2s, providing a valuable approach for visualizing distinct BM ILC2 subsets.

We next examined the cellular composition of the ID2^hi^CXCR6⁻ and ID2^hi^CXCR6^hi^ compartments to determine the extent to which other immune populations shared these reporter profiles (**Figure 1C-E**). Within the Lin^-^ID2^hi^CXCR6^⁻^ fraction, NK cells (CD3ε⁻NK1.1⁺) represented the predominant population (53 ± 23%), followed by ILC/NK progenitors (20.9 ± 7.9%) and ST2⁺ ILC2s (10.3 ± 7.2%). NKT cells accounted for 8.2 ± 7.3%, CD4⁺ T cells for 3 ± 1.2%, and CD8⁺ T cells for 1 ± 0.6% of this compartment (**Figure 1E**). NK cell identity was further supported by NKp46 expression, while NKT cells comprised both CD4⁺ and CD4⁻ subsets (**Figure S1A**).

In contrast, the Lin⁻ID2^hi^CXCR6^hi^ compartment was predominantly composed of NKT cells (40 ± 12.9%), together with CD4⁺ T cells (15.5 ± 5.4%) and CD8⁺ T cells (3.4 ± 1.6%). ILC2s represented 7.2 ± 1.9% of this compartment, comprising both ST2⁻ (approximately 3%) and ST2⁺ (approximately 4%) subsets. NK cells (5.8 ± 3.4%) and ILC/NK progenitors (2.9 ± 3.2%) were less abundant (**Figure 1E**). NKp46 expression confirmed the identification of NK cells, and NKT cells again included both CD4⁺ and CD4^⁻^ subsets (**Figure S1B**).

We also examined the ID2^med^ compartment, which contained no detectable ILC2s and was predominantly composed of Lin⁺ myeloid cells. Among the lymphoid populations identified in this compartment, CD8⁺ T cells were the most abundant (**Figure S2A, B**).

Together, these results demonstrate that although ID2 and CXCR6 reporters label all ST2⁻ ILC2s and a substantial proportion of ST2⁺ ILC2s, these reporter combinations are also shared by several other BM lymphoid populations, particularly NK, NKT and T cells. Therefore, additional surface markers are required to specifically distinguish ILC2s from other ID2 and CXCR6 expressing cells for their identification by confocal microscopy.

### Surface marker profiling identifies candidate markers for distinguishing BM ILC2s from other ID2 expressing populations

To define candidate surface markers for distinguishing BM ILC2 from other reporter-positive cells in tissue sections, we used spectral flow cytometry to compare the expression of selected surface markers across the different BM lymphoid populations identified in Figure 1. As expected, antibodies against CD4, CD8, and NK1.1 allowed the discrimination of T and NK cell populations within the ID2^hi^CXCR6⁻ and CXCR6^+^ gates (**Figure 2A**).

**Figure 2.**
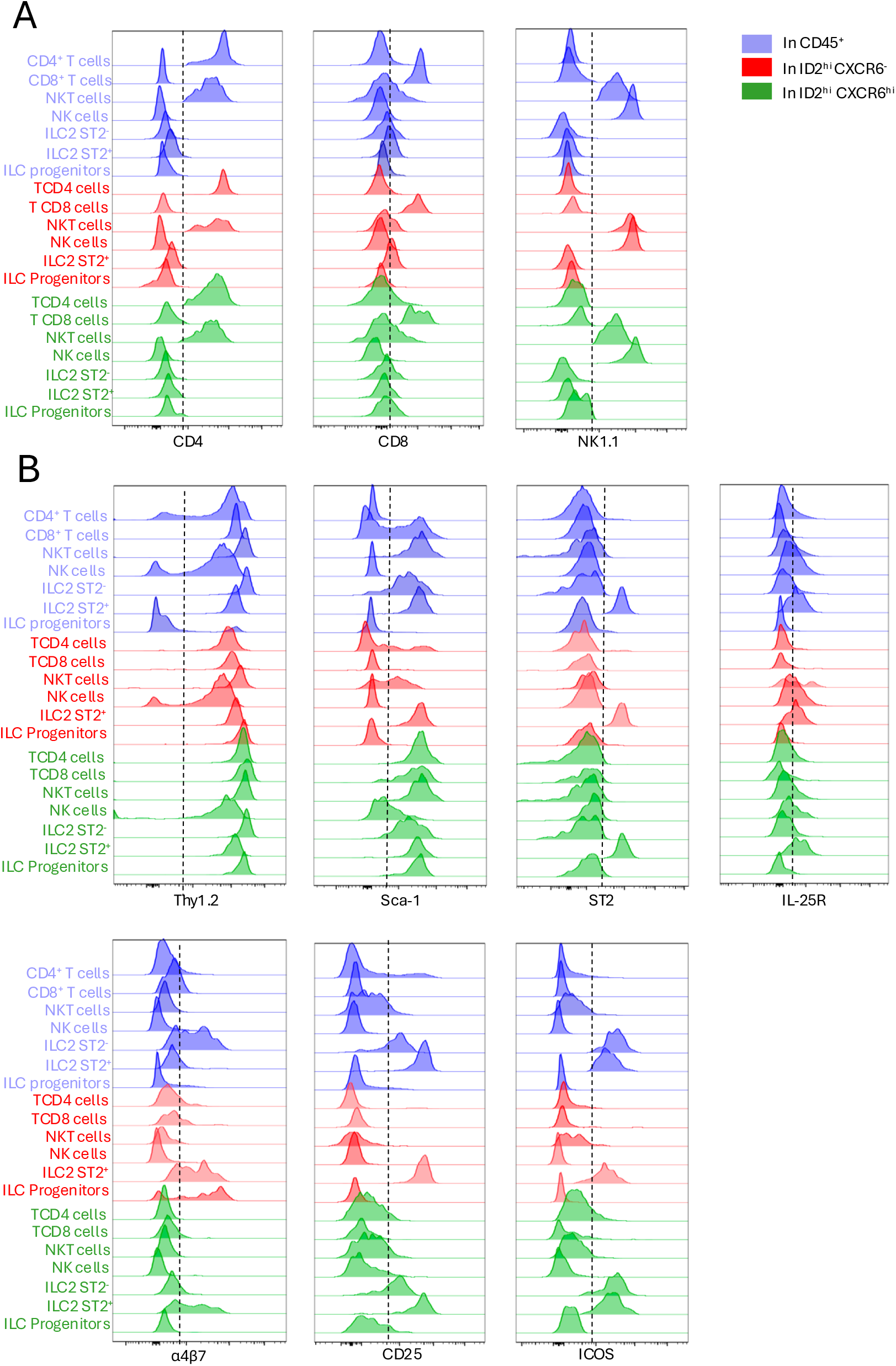
Surface marker profiling identifies candidate markers for distinguishing BM ILC2s from other ID2-expressing populations. (A, B) Overlay histograms showing expression of the indicated surface markers across gated BM lymphoid populations, analyzed within total CD45⁺ cells (blue), ID2ʰⁱCXCR6⁻ cells (red), or ID2ʰⁱCXCR6ʰⁱ cells (green). Markers include CD4, CD8, NK1.1, Thy1.2, Sca-1, ST2, IL-25R, α4β7, CD25 and ICOS.

Thy1.2 was broadly expressed on mature lymphocytes at varying levels, and Sca-1 was expressed by all populations within the ID2^hi^CXCR6^hi^ fraction (**Figure 2B**). The receptor ST2 was restricted to the ST2⁺ ILC2 subset among the populations examined. IL-25R showed a similar profile although it was also detected on a minor subset of NK cells (**Figure 2B**). The integrin α4β7 was expressed by ST2^+^ ILC2s but was also detected on ILC progenitors within the ID2^+^CXCR6⁻ compartment. Finally, CD25 was expressed by both ST2⁻ and ST2⁺ ILC2 subsets, whereas ICOS was highly expressed by ILC2s while lower levels of expression were also detected in a small proportion of NK, NKT and T cells within both ID2^hi^CXCR6⁻ and ID2^hi^ CXCR6^hi^ compartments (**Figure 2B**).

### Evaluation of surface marker combinations for the identification of BM ILC2s

We next evaluated whether combining CD25 and ICOS could improve the identification of ST2⁻ and ST2⁺ ILC2 subsets within the ID2⁺ CXCR6^hi^ and ID2⁺ CXCR6⁻ compartments. We assessed the composition of the CD25⁺ICOS⁺ population to determine the proportion of ILC2s and other lymphoid cells within this selected compartment. The inclusion of ST2 as an additional marker allowed for the specific delineation of ST2⁺ ILC2s, as the ST2⁺ fraction showed no detectable contamination by NKT, NK, or T (CD4/CD8) cells (**Figure 3A-C**). In contrast, within the selected ST2⁻ fraction, the ST2⁻ subset exhibited low levels of contamination by other lymphoid populations, comprising NKT cells (6,6 ± 0,1 %), NK cells (0,9 ± 0,5 %), CD4⁺ (1,6 ± 0,3 %) and CD8⁺ T cells (0,3 ± 0,3 %) consistent with the low levels of ICOS expression observed in a small proportion of cells within these populations (**Figure 2B**). Thus, CD25 and ICOS staining enabled the selection of a population highly enriched in ILC2s, with approximately 90% purity within the ID2^hi^CXCR6^hi^ compartment. The combination of CD25/ICOS with ST2 allows reliable identification of ST2⁺ ILC2s by flow cytometry within ID2^hi^ CXCR6^hi^ and ID2^hi^CXCR6⁻ fractions. In contrast, the ST2⁻ subset identified using CD25 and ICOS retains a small proportion of contaminating lymphoid cells. Therefore, additional exclusion markers, including CD3, NK1.1 or NKp46 may be required to distinguish ILC2s from these populations when developing an imaging strategy.

**Figure 3.**
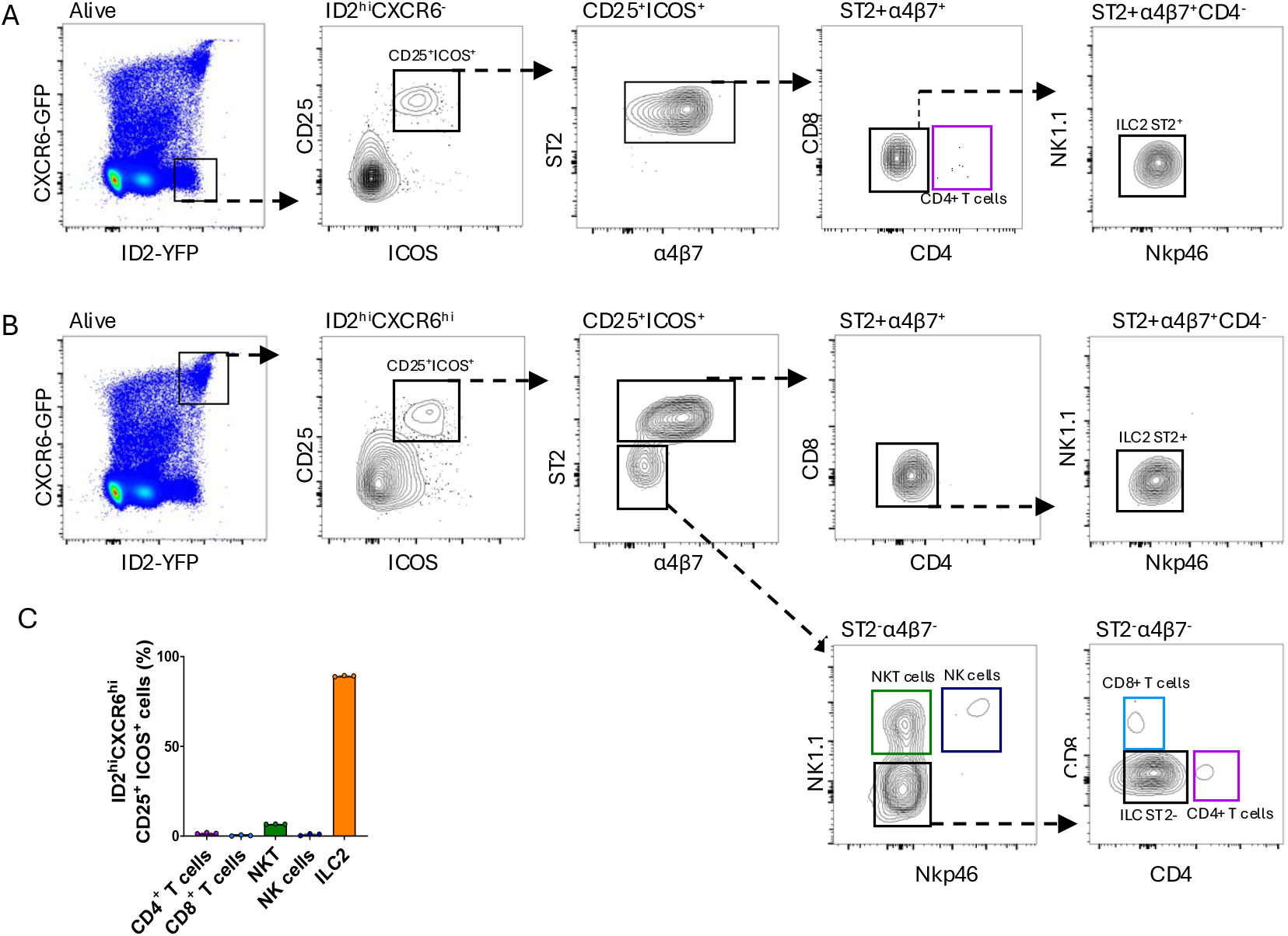
Evaluation of surface marker combinations for the identification of BM ILC2s. (A, B) Representative flow cytometry plots showing the sequential gating strategy based on CD25, ICOS, ST2 and α4β7 expression within the (A) ID2^hi^CXCR6⁻ and (B) ID2^hi^CXCR6^hi^ BM compartments. ST2⁺α4β7⁺ cells were further characterized by CD4, CD8, NK1.1 and NKp46 expression. Within the ID2^hi^CXCR6^hi^ compartment, ST2⁻α4β7⁻ cells were further analyzed to distinguish NK, NKT, CD4⁺ T, CD8⁺ T and ST2^-^ ILC2 populations. (C) Quantification of the indicated lymphoid populations among ID2^hi^CXCR6^hi^CD25⁺ICOS⁺ cells. Data are presented as mean ± SD. Data were obtained from two independent experiments involving a total of seven mice.

### Molecular characterization of BM ILC2s and other Id2 and CXCR6 expressing lymphoid populations

To further characterize the lymphoid populations expressing ID2 and CXCR6 in the BM lymphoid, ILC2s, T/NKT cells and NK cells were sorted by flow cytometry (**Figure S3**) and analyzed using single-cell RNA sequencing (scRNA-seq). The UMAP revealed 18 lymphoid clusters in the BM (excluding B cells), which could be identified based on the expression of specific marker genes (**Figure 4A-B** and Material and Methods). A small population of cycling cells was identified among immature NKT cells (**Figure 4C**). As observed by spectral flow cytometry, cells expressing high levels of *Id2* transcripts were predominantly associated with mature NK cells and their precursors (**Figure 1E**, **Figure 4B, D**). Cells co-expressing *Id2* and *Cxcr6* included ILC2 as well as immature NK cells, NKT2 cells, CD8 resident memory T cells (CD8 Trm) and γδTh17 cells confirming that co-expression of these genes is not restricted to ILC2s. However, unlike our observations by spectral flow cytometry, a distinct population of *Id2*^+^*Cxcr6*^+^ CD4^+^ T cells was not readily identifiable in the initial clustering analysis. Since these cells may represent a rare subset in the BM, subclustering was performed on the clusters corresponding to CD4^+^ T cells (Treg and naive T cells) as well as the nearest clusters in the UMAP (CD8 Trm, CD8 Tcm and Vγ1^+^ γδT cells). This revealed a rare CD4^+^ T cell subset that was distinct from naïve and regulatory T cells (Cd44^+^, Sell⁻, Foxp3⁻) and expressed both *Id2* and *Cxcr6* (**Figure S4**; cluster 6). These cells shared transcriptional features with previously described CCL5^+^ cytotoxic CD4^+^ T cells that accumulate in the BM with age [31]. Several genes expressed by ILC2s helped discriminate them from other lymphoid populations co-expressing *Id2* and *Cxcr6*, including *Gata3*, *Il1rl1*, *Il17rb* and *Stab2* (**Figure 4B**). In addition, ILC2s lacked detectable *Cd5* and *Ncr1* transcripts, whereas these genes were expressed by T/NKT and NK cell populations respectively.

**Figure 4.**
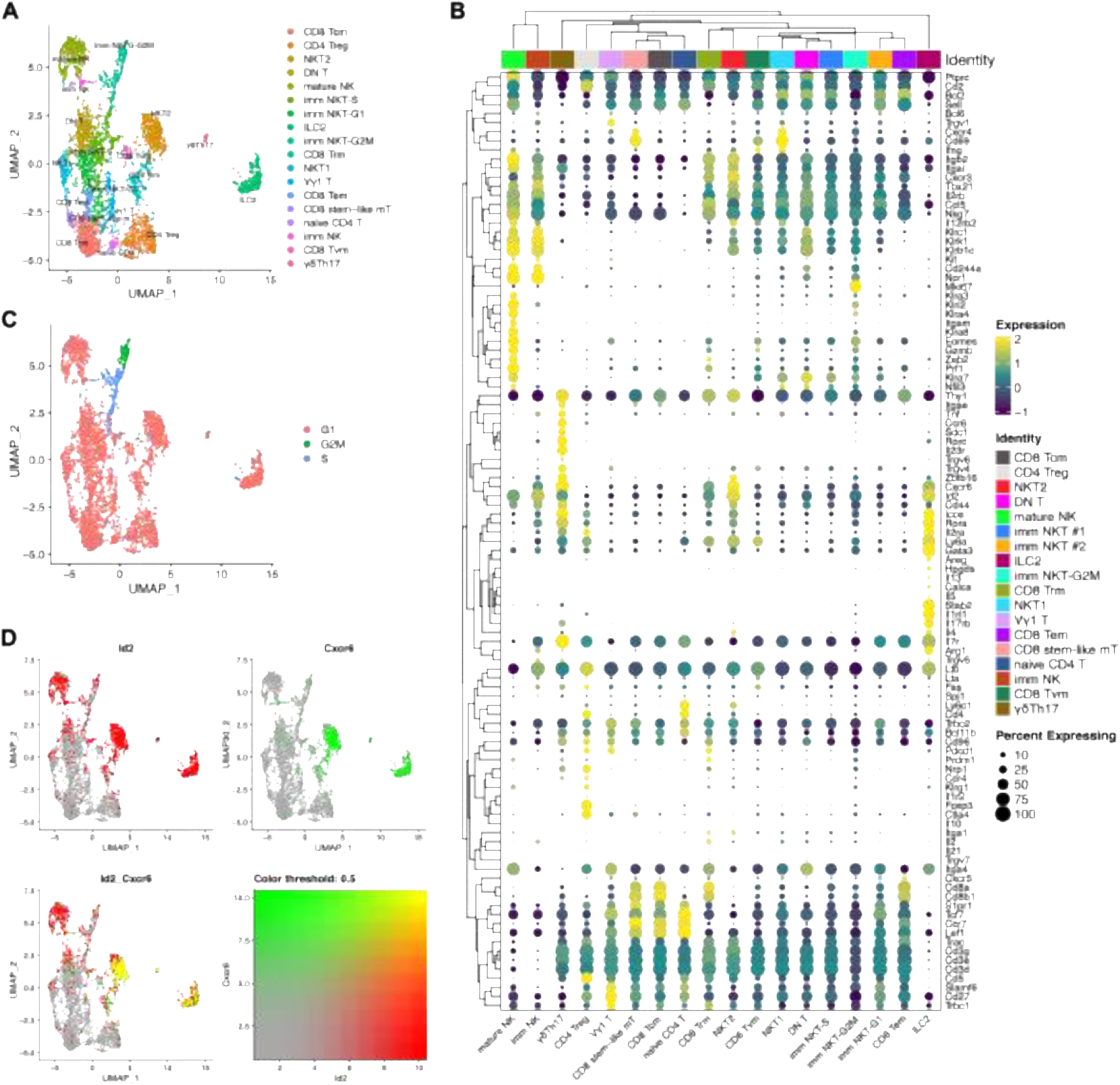
Single-cell transcriptomic profiling reveals the diversity of Id2- and Cxcr6-expressing BM lymphoid populations. Lymphoid cells from murine BM were sorted by flow cytometry and analyzed by single-cell RNA sequencing (scRNA-seq). (A) UMAP representation of the identified T, NK and NKT cell subsets. (B) Expression patterns of selected lymphoid marker genes across clusters. (C) Cell-cycle assignments for cells represented on the UMAP. (D) Co-expression of Id2 and Cxcr6 transcripts across the analyzed cells.

Overall, these results confirmed the presence of several lymphoid subsets expressing both ID2 and CXCR6 in the BM, highlighting the need for additional markers to distinguish ILC2s from other double positive cells. Furthermore, the differential expression of *Cd5* and *Ncr1* provided a rationale for using CD5 and Nkp46 as exclusion markers in subsequent confocal imaging experiments.

### Confocal imaging reveals the spatial distribution of putative ILC2 within the BM

To visualize BM ILC2s in situ using confocal microscopy, we combined the Id2-YFP and Cxcr6-GFP reporters with additional surface markers to distinguish ILC2s from other reporter-positive lymphoid populations. Although ST2 and ICOS were identified as useful markers for ILC2 identification by flow cytometry, neither could be reliably detected by immunofluorescence under the bone processing and staining conditions used in this study. We therefore adopted an alternative approach, excluding T and NKT cells based on CD5 expression and NK cells based on NKp46 expression in the BM of ID2^YFP/+^CXCR6^GFP/+^ reporter mice. YFP^+^ and YFP^+^GFP^+^ cells were found throughout the BM (**Figure 5A**). Furthermore, these cells were either isolated or organized into multicellular clusters (**Figure 5B**). CD5^+^ T/NKT cells were detected among the YFP^+^GFP^+^ cells while NKp46^+^ NK cells were also identified within this double positive compartment (**Figure 5C**). A subset of YFP^+^GFP^+^CD5⁻NKp46⁻ cells also expressed CD90, providing additional support for their identification as putative ILC2s (**Figure S5**).

**Figure 5.**
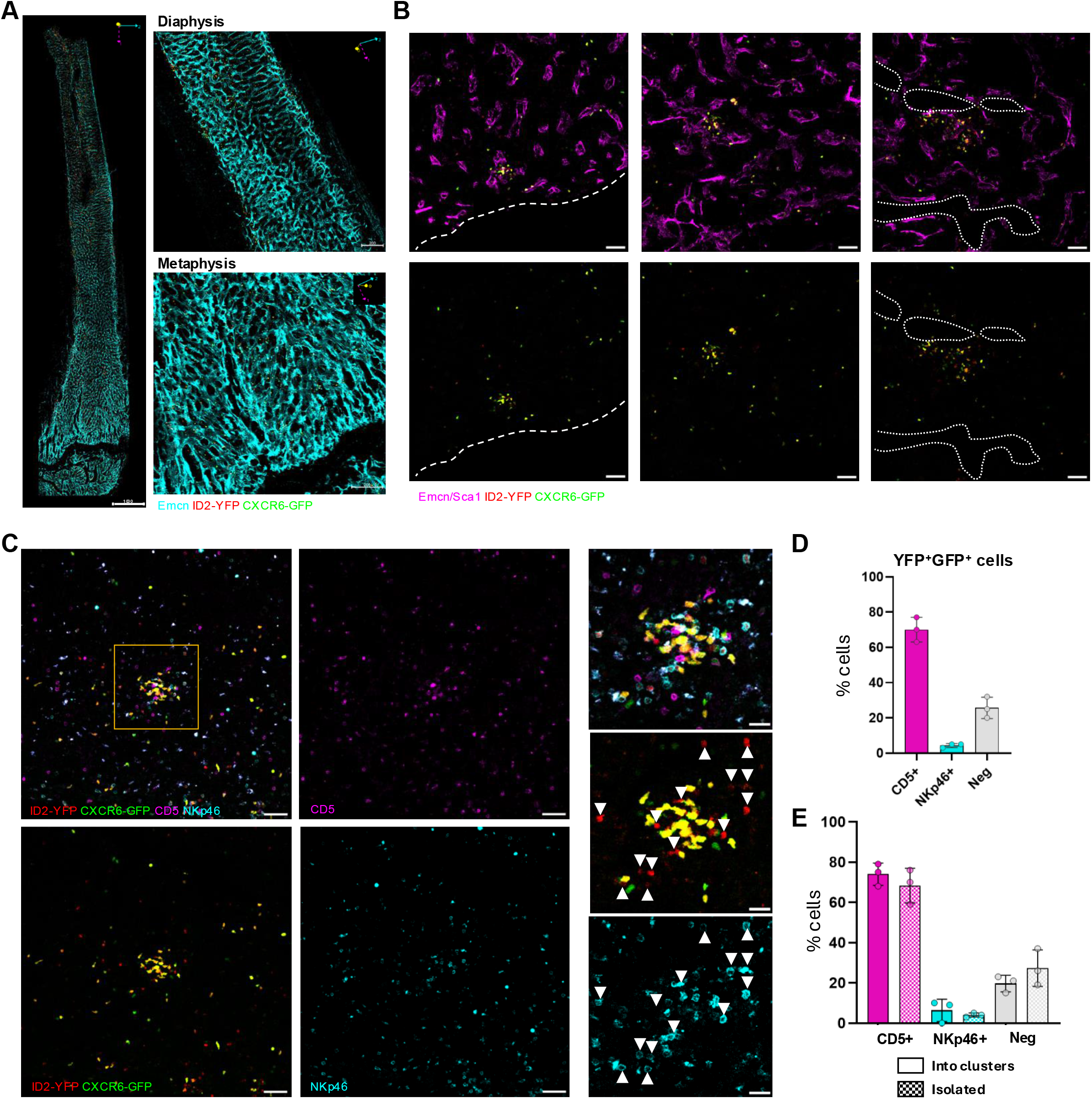
Confocal imaging reveals the spatial distribution of putative ILC2s within the bone marrow. (A) Whole-mount confocal imaging of a femur stained for endomucin (EMCN) to visualize sinusoidal endothelium, together with YFP and GFP reporter fluorescence. Images were acquired using a 20× objective (left panel); enlarged views of the diaphysis and metaphysis are shown (right panels). The x, y and z orientations are indicated. Scale bars: 1,000 µm (left); 200 µm (right). (B) BM sections stained for EMCN and Sca-1 (detected in the same fluorescence channel), showing YFP and GFP reporter fluorescence. Dotted lines indicate the bone surface. Scale bars: 50 µm. (C) BM sections showing YFP and GFP reporter fluorescence together with CD5 and NKp46 staining. Enlarged views of the outlined region are shown. Arrows indicate YFP^+^ cells. Scale bars: 50 µm (left and central panels); 20 µm (right panels). (D) Proportions of CD5⁺, NKp46⁺ and CD5^-^NKp46^-^ (Neg) cells among YFP⁺GFP⁺ cells identified by confocal microscopy (n = 3 mice). No CD5⁺NKp46⁺ double-positive cells were observed. (E) Proportions of these populations among YFP⁺GFP⁺ cells observed as isolated cells or within multicellular clusters, as defined in the Materials and Methods. For panels D and E, each dot represents one individual mouse (n = 3 mice).

Quantification of the different YFP^+^GFP^+^ cell populations revealed that CD5^+^ T/NK T cells were the most abundant, although NKp46^+^ NK cells and CD5⁻NKp46⁻ including putative ILC2s were also present (**Figure 5D**). These different lymphoid populations were observed either as isolated cells or within multicellular clusters, some of which contained cells belonging to distinct lymphoid populations (**Figures 5C, E**). Altogether, these results demonstrate that ID2^YFP/+^ CXCR6^GFP/+^ mice combined with additional surface markers provide a useful approach for visualizing putative BM ILC2s and examining their spatial distribution in situ.

### Flow cytometric validation of ILC2 enrichment among reporter-positive BM cells

Although the combination of ID2 and CXCR6 reporters with CD5 and NKp46 staining allowed us to identify cells displaying an ILC2 compatible phenotype in BM sections, this approach could not formally exclude the presence of NK/ILC progenitors sharing a similar phenotype. To assess their relative contribution, we performed additional flow cytometric analyses of ID2^hi^CXCR6^hi^ BM cells (**Figure 6A**). Following exclusion of Lin^+^ cells and CD3ε⁺, NKp46⁺ and NK1.1⁺ populations, we examined the remaining Thy1⁺ cells. ILC2s, identified by ICOS and IL7Rα expression, accounted for 93.9 ± 2.5% of the selected population, whereas ICOS⁻IL7Rα^low/+^ cells represented 5.3 ± 2.0% (mean ± SD, n = 4 mice) (**Figure 6B**). The phenotype of this minor population was compatible with that of putative ILC progenitors. These results demonstrate that ILC2s constitute the predominant population within this phenotypically defined compartment, providing additional support for the identification of reporter-positive cells observed by confocal microscopy as putative ILC2s.

**Figure 6.**
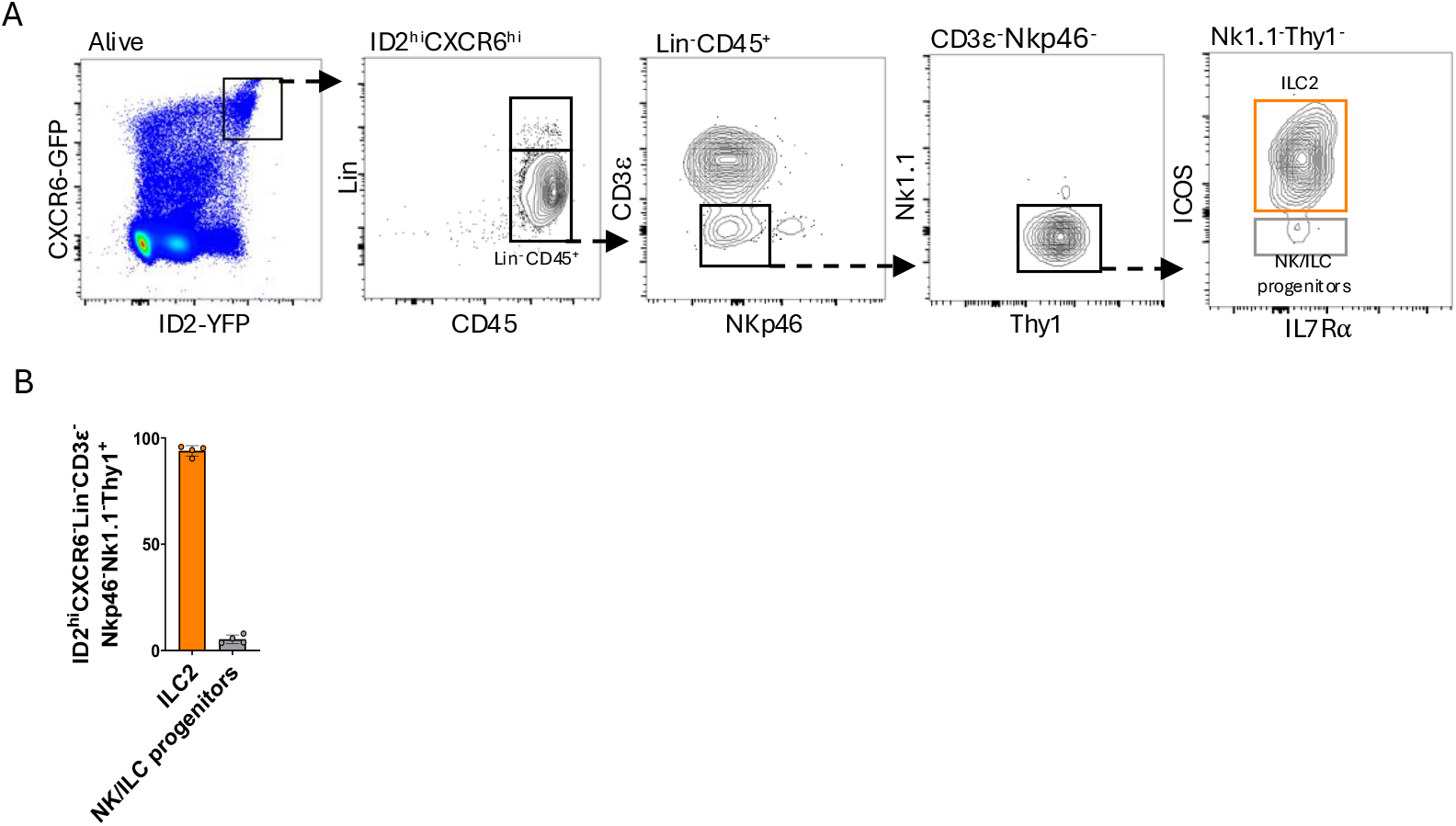
Flow cytometric analysis confirms the predominance of ILC2s among ID2^hi^CXCR6^hi^ BM cells after exclusion of major T and NK cell populations. (A) Representative flow cytometry plots showing the gating strategy used to identify ILC2s and a minor population compatible with putative NK/ILC progenitors among ID2^hi^CXCR6^hi^ BM cells from ID2/CXCR6 dual-reporter mice. Live ID2^hi^CXCR6^hi^ cells were selected, followed by gating on Lin⁻CD45⁺ cells and exclusion of CD3ε⁺, NKp46⁺ and NK1.1⁺ cells. Among the remaining Thy1⁺ cells, ILC2s were identified as ICOS⁺IL7Rα⁺ cells, whereas the minor ICOS⁻IL7Rα^low/+^ population was phenotypically compatible with putative NK/ILC progenitors. (B) Proportions of ILC2s and ICOS⁻IL7Rα^low/+^ cells (putative ILC progenitors) among ID2^hi^CXCR6^hi^ Lin⁻CD3ε⁻NKp46⁻NK1.1⁻Thy1⁺ BM cells. Each dot represents one mouse (n = 4). Data are presented as mean ± SD.

## Discussion

The identification and localization of ILCs within tissues remain challenging, particularly in the BM, where ILC2s represent a minor population among heterogeneous lymphoid cells. Although reporter mice have considerably advanced our understanding of ILC development and heterogeneity, their use for identifying mature ILC subsets in situ requires careful consideration of reporter expression in other lymphoid populations. In this study, we evaluated the use of *Id2*/*Cxcr6* dual-reporter mice to identify and visualize BM ILC2s. By combining spectral flow cytometry, scRNA-seq and confocal microscopy, we characterized the diversity of lymphoid populations expressing ID2 and CXCR6 and identified complementary markers to improve the discrimination of ILC2s from other reporter-positive cells. Our findings highlight both the potential and the limitations of this reporter combination for studying the spatial distribution of BM ILC2s.

ID2 is an essential regulator of ILC development, and *Id2* reporter mice have been instrumental in identifying BM ILC progenitors and investigating their differentiation potential [10,20,19]. However, previous studies have established that *Id2* expression is not restricted to helper-like ILCs but is also detected in NK cells and their progenitors [20]. CXCR6 was initially identified as a marker associated with ILC development, with its expression marking a progressive restriction of lymphoid progenitor potential toward innate lymphoid lineages [6]. Subsequent studies further characterized CXCR6⁺α4β7⁺ BM progenitors and demonstrated their differentiation potential toward multiple ILC lineages [8]. Notably, previous analyses using *Id2*/*Cxcr6* dual-reporter mice demonstrated heterogeneous CXCR6 expression among BM ILC precursors and ILC2 progenitors [28].

Consistent with these observations, our spectral flow cytometry analyses revealed that ID2 expressing BM cells comprise a heterogeneous mixture of NK cells, NKT cells, conventional T cells, ILC progenitors and ILC2s. Importantly, although both ST2⁻ and ST2⁺ ILC2 subsets expressed high levels of ID2, they differed in CXCR6 expression. ST2⁻ ILC2s were uniformly CXCR6⁺, whereas ST2⁺ ILC2s comprised both CXCR6⁺ and CXCR6⁻ populations. Thus, while combining *Id2* and *Cxcr6* reporters facilitates the detection of BM ILC2s, their co-expression is neither exclusive to ILC2s nor shared by all BM ILC2 subsets. Consequently, restricting imaging analyses to double-positive cells would exclude a fraction of the ST2⁺ ILC2 population.

Our study also emphasizes the importance of considering both the expression levels and the specificity of surface markers when identifying ILC2s among other BM lymphoid populations. Consistent with previous studies reporting high ICOS expression by ILC2s [32,33], CD25 and ICOS were useful for identifying both ST2⁻ and ST2⁺ ILC2 subsets. ICOS was expressed at high levels by ILC2s, whereas low levels of expression were detected in small fractions of NK, NKT and T cells. Importantly, combining high ID2 and CXCR6 reporter expression with CD25 and ICOS staining resulted in substantial enrichment of ILC2s, which represented approximately 90% of the ID2^hi^CXCR6^hi^CD25⁺ICOS⁺ population. Nevertheless, this population still contained a small proportion of other lymphoid cells, predominantly NKT cells, indicating that this combination yields a highly enriched ILC2 population without achieving complete specificity. The additional use of ST2 allowed a more selective identification of ST2⁺ ILC2s, whereas the identification of ST2⁻ ILC2s remained more challenging because of their phenotypic overlap with other lymphoid populations. However, despite their usefulness in flow cytometry, neither ICOS nor ST2 could be reliably detected by immunofluorescence under our bone processing and staining conditions, limiting their application to the identification of ILC2s in BM sections. These observations illustrate the limitations of relying on a restricted number of positive markers and support the use of complementary exclusion markers to improve the identification of BM ILC2s.

Single-cell transcriptomic analysis provided an additional level of characterization by revealing the diversity of lymphoid populations co-expressing *Id2* and *Cxcr6*. Besides ILC2s, these included NK cells, NKT2 cells, tissue-resident memory CD8⁺ T cells and γδ T cells. The identification of a rare population of CD4⁺ T cells expressing both genes further illustrates the complexity of the BM lymphoid compartment. These cells shared transcriptional features with cytotoxic CD4⁺ T cells recently described in the BM [31], although their precise relationship to this population remains to be established. Conversely, the ILC2 population was characterized by the expression of genes associated with its identity, including *Gata3*, *Il1rl1* and *Il17rb*, together with the absence of detectable *Cd5* and *Ncr1* transcripts. These observations provided a rationale for using CD5 and NKp46 as exclusion markers in subsequent imaging experiments.

An objective of our study was to translate the phenotypic information obtained by flow cytometry into a strategy suitable for confocal imaging of BM sections. This represents a particular challenge because markers that reliably distinguish lymphoid populations in cell suspensions are not necessarily compatible with tissue processing and immunofluorescence staining. We therefore combined the *Id2* and *Cxcr6* reporters with CD5 and NKp46 staining to exclude major T/NKT and NK cell populations. This approach allowed us to visualize YFP⁺GFP⁺CD5⁻NKp46⁻ cells, some of which also expressed CD90, consistent with an ILC2 phenotype. To further evaluate the contribution of ILC2s and NK/ILC progenitors to this reporter-positive compartment, we performed complementary flow cytometric analyses. These analyses revealed that ILC2s accounted for more than 90% of the selected population, whereas ILC/NK progenitors represented only a minor fraction. This strong enrichment provides additional support for the identification of the cells observed by confocal microscopy as putative ILC2s.

Confocal imaging further revealed that reporter-positive lymphoid cells were distributed throughout the BM, either as isolated cells or within multicellular clusters. These observations provide initial information on the spatial organization of lymphoid populations expressing ID2 and CXCR6 in the BM. The presence of distinct lymphoid subsets within the same clusters suggests that these structures are not restricted to a single cell population. However, their cellular composition, spatial relationships with stromal and vascular compartments, and potential functional significance remain to be determined. In particular, the observation of clustered cells does not establish whether they engage in direct interactions or participate in the organization of specific BM microenvironments.

A limitation of our imaging strategy is that it does not provide an exhaustive or exclusive identification of BM ILC2s. Heterogeneous CXCR6 expression excludes a fraction of ST2⁺ ILC2s from the double reporter-positive population, while the exclusion of CD5⁺ and NKp46⁺ cells does not formally eliminate NK/ILC progenitors or other rare populations sharing a similar phenotype. Although CD90 expression provides additional phenotypic support, it cannot establish ILC2 identity on its own. Moreover, because the gating strategies used for flow cytometry and confocal imaging are not strictly identical, the precise proportion of ILC2s among the cells visualized in situ cannot be determined.

Overall, our study establishes a complementary flow cytometric, transcriptomic and imaging framework for characterizing BM ILC2s using *Id2*/*Cxcr6* dual-reporter mice. The combination of reporter expression and complementary surface markers enables substantial enrichment of ILC2s by flow cytometry, while the identification of reporter-positive cells in BM sections is supported by their phenotypic characterization and the predominance of ILC2s within a closely related population analyzed by flow cytometry. This approach provides a basis for investigating the spatial distribution of putative BM ILC2s and their relationships with other cellular components of the hematopoietic microenvironment.

## ACKNOWLEDGMENTS

We acknowledge the Cytometry and Biomarkers Unit of Technology and Service (CB UtechS) at Institut Pasteur, the flow cytometry platform and the MRic microscopy platform of the UAR Biosit in Rennes for support in conducting this study. We thank the staff of the animal facility of the Institut Pasteur for mouse care. We thank the core bioinformatics platform of the INSERM U1236 for their expert support with scRNA-seq data analysis. This work has been supported by Institut Pasteur, Université Paris Cité, CNRS and by grants from INSERM by the French Government (National Research Agency) ANR project CINIMA (#ANR-22-CE15-0005). C.L. was was financially supported by the French National Research Agency (#ANR-22-CE15-0005) and by the Société Française d’Hématologie (SFH). P.H. was financially supported by the French National Research Agency (#ANR-22-CE15-0005) and by the Labex Revive. R.G. and G. J. acknowledge salary support from the Université Paris Cité. EGP acknowledges salary support from Institut Pasteur.

## AUTHOR CONTRIBUTIONS

C.L. performed scRNA-seq library analysis, confocal imaging, analyzed data, constructed figures, and participated to the manuscript writing. P.H. followed the breeding of animals, performed spectral flow cytometry experiments analyzed data, constructed figures, and participates to the manuscript writing. G.J. performed spectral flow cytometry experiments analyzed data. E.G.M. Contributed to resources and scientific discussions. R.G. and S.J.C.M directed the research, designed the studies, analyzed data, secured funding and wrote the manuscript.

## DECLARATION OF INTERESTS

The authors declare no competing interests.

## Abbreviations

Type 2 innate lymphoid cells (ILC2s); bone marrow (BM); single-cell RNA sequencing (scRNA-seq).

---

**Figure S1.**
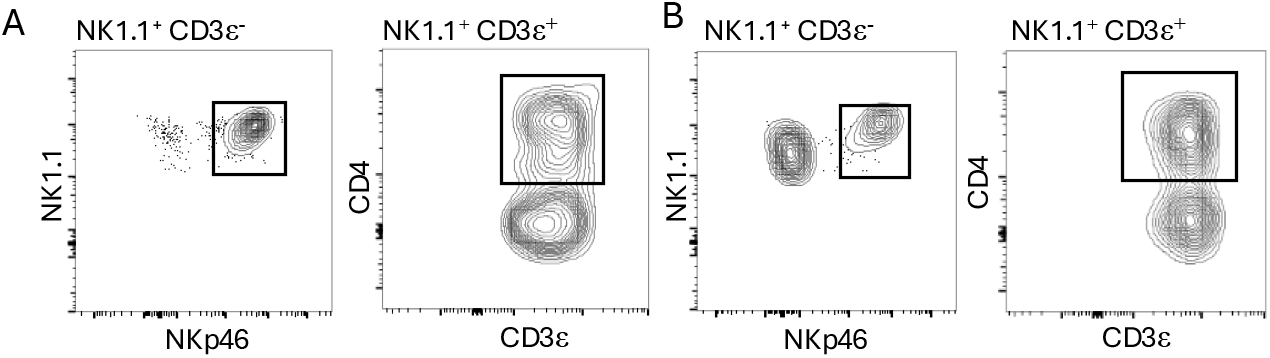
Characterization of NK and NKT cell populations within ID2ʰⁱCXCR6^-^ and ID2ʰⁱCXCR6ʰⁱ BM compartments. Representative flow cytometry plots showing NK and NKT cell populations within (A) ID2ʰⁱCXCR6^-^ and (B) ID2ʰⁱCXCR6ʰⁱ gates.

**Figure S2.**
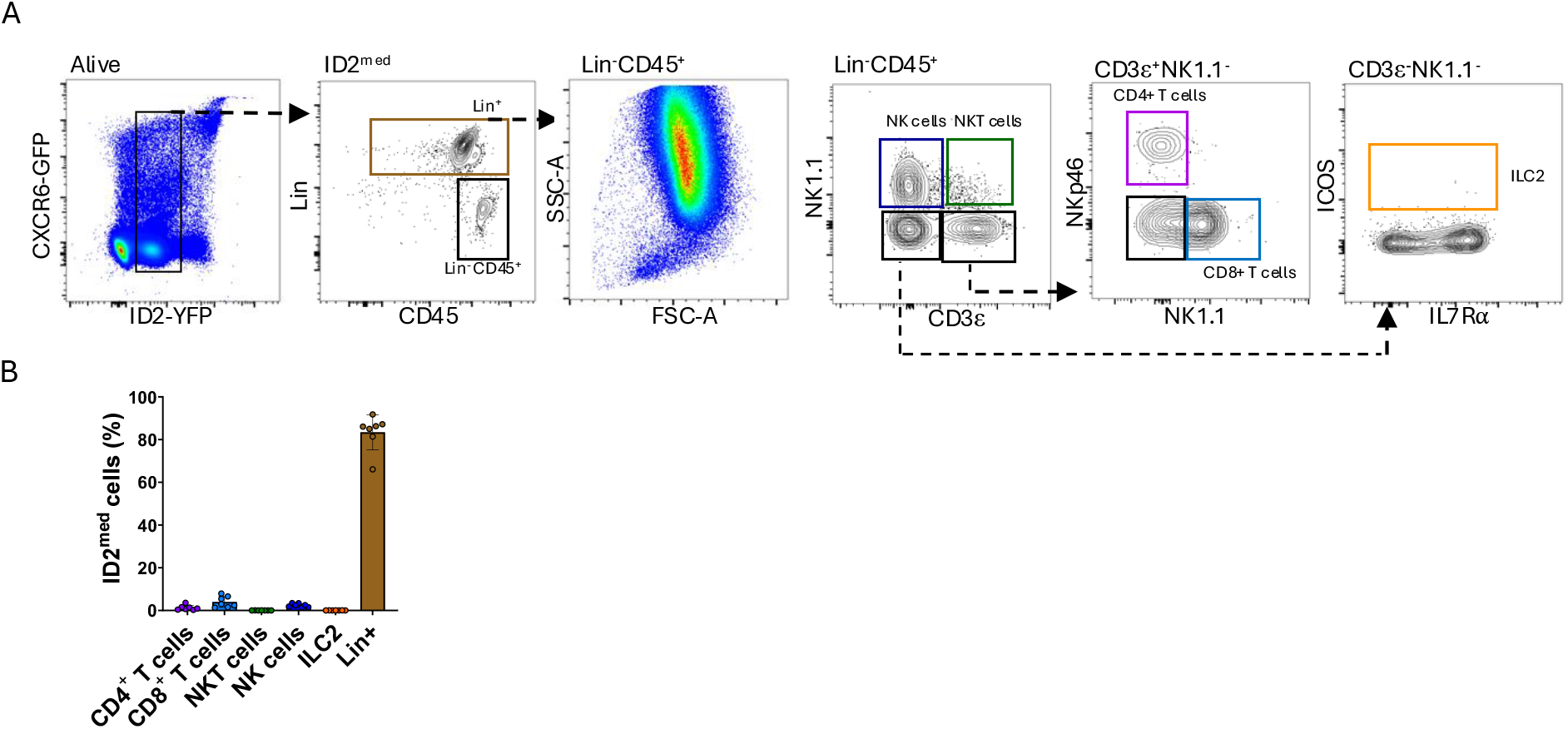
The ID2ᵐᵉᵈ BM compartment is enriched in lineage-positive cells and lacks detectable ILC2s. (A) Representative flow cytometry plots showing the distribution of Lin⁺ cells (CD11b, Gr-1, CD19, Ter119), T cells, NKT cells and NK cells within the ID2ᵐᵉᵈ compartment. (B) Quantification of the indicated populations among total ID2ᵐᵉᵈ cells. Data are presented as mean ± SD. Data were obtained from two independent experiments involving a total of seven mice.

**Figure S3.**
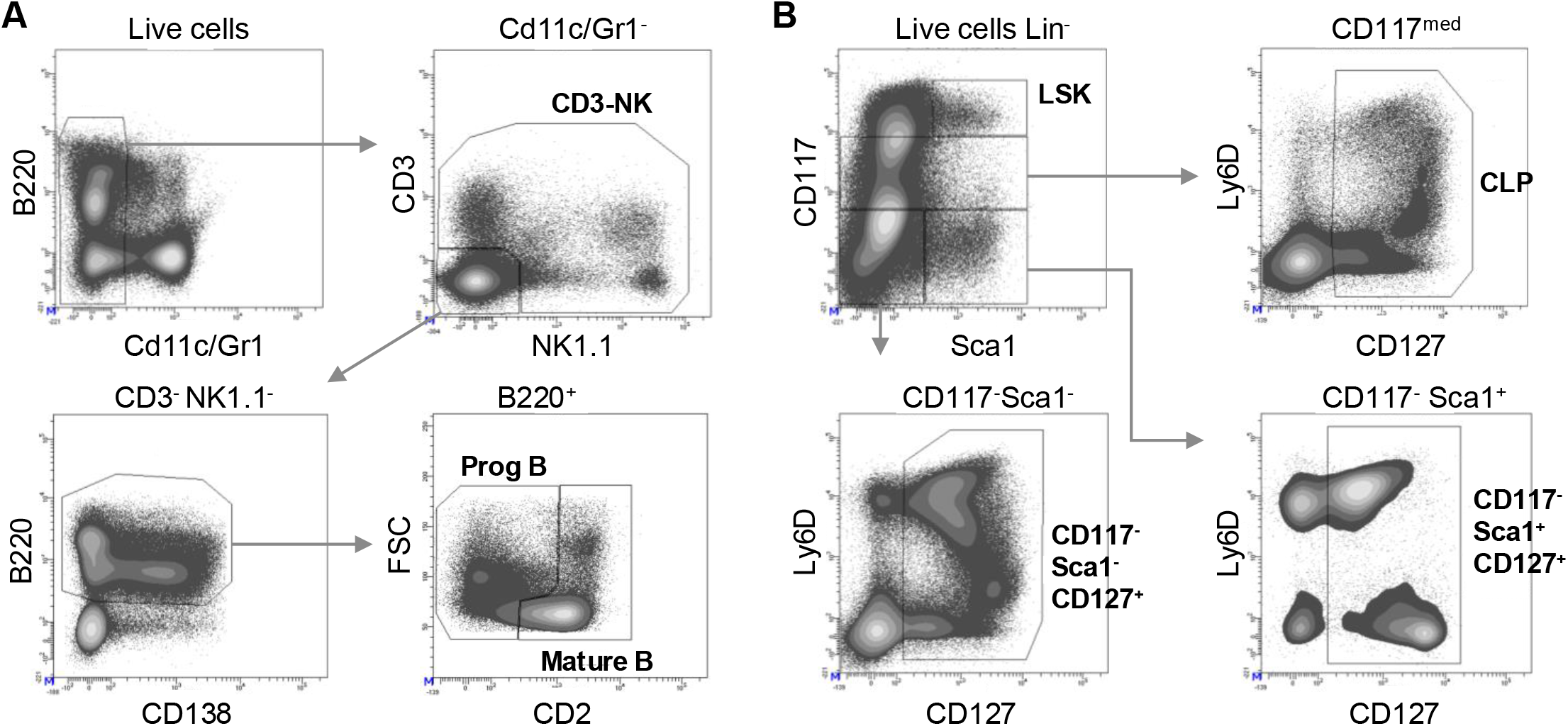
Flow cytometric sorting strategy for single-cell RNA sequencing of BM lymphoid populations. BM cells were flushed and stained with the antibodies indicated in the panels and described in Materials and Methods. (A) Gating strategy for lineage-marker-expressing lymphoid populations. (B) Gating strategy for lymphoid populations within the lineage-negative BM fraction, including ILC2s.

**Figure S4.**
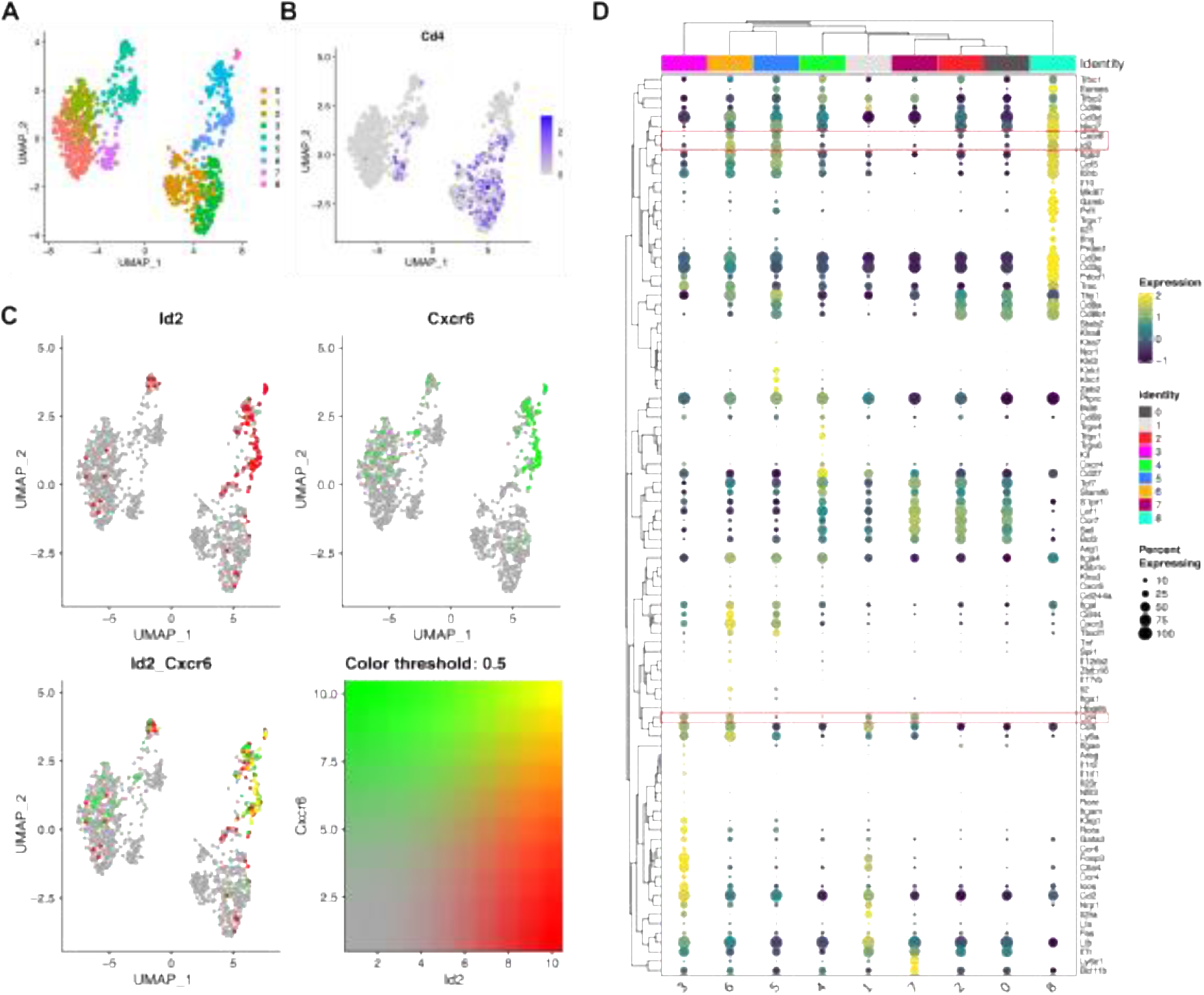
Subclustering reveals a rare Id2- and Cxcr6-expressing CD4⁺ T-cell population in BM. Clusters annotated as regulatory T cells, naïve T cells, CD8 tissue-resident memory T cells, CD8 central memory T cells and Vγ1⁺ γδ T cells were extracted from the scRNA-seq dataset shown in Figure 4 for refined clustering. (A) UMAP representation of the subclustered populations. (B) Cd4 expression across the UMAP. (C) Co-expression of Id2 and Cxcr6 transcripts. (D) Expression patterns of selected lymphoid marker genes across clusters. Red boxes highlight Id2, Cxcr6 and Cd4 expression.

**Figure S5.**
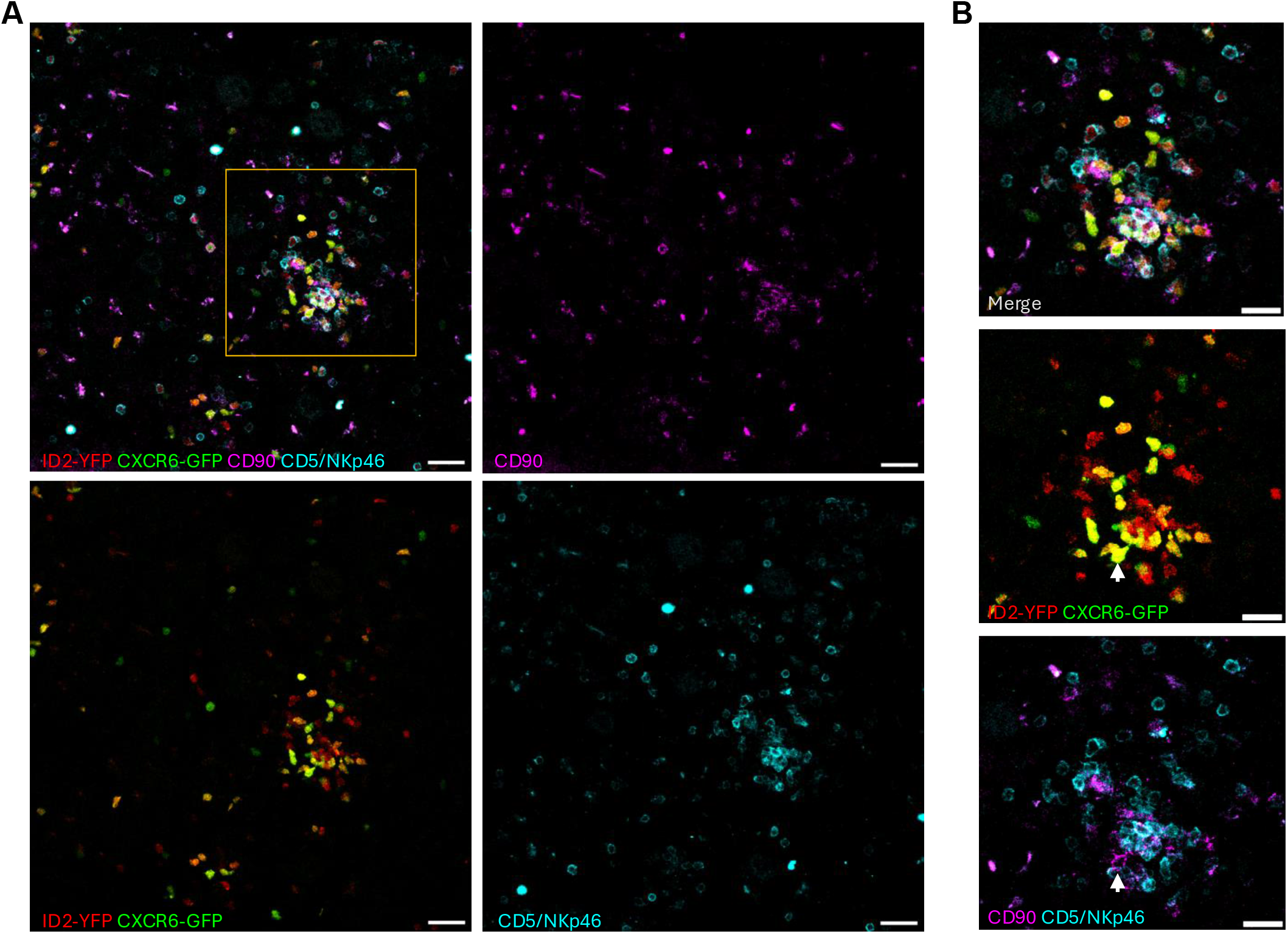
CD90 expression by YFP⁺GFP⁺CD5⁻NKp46⁻ cells in BM sections. (A) BM sections stained for CD5 and NKp46 (detected in the same fluorescence channel) and CD90, together with YFP and GFP reporter fluorescence. Scale bars: 30 µm. (B) Enlargement of the outlined region in the upper left panel of (A). The arrow indicates a YFP⁺GFP⁺CD5⁻NKp46⁻CD90⁺ cell. Scale bars: 20 µm.

## Notes

### Competing Interest Statement

The authors have declared no competing interest.

## References

[1] G. Gasteiger, X. Fan, S. Dikiy, S.Y. Lee, A.Y. Rudensky, Tissue residency of innate lymphoid cells in lymphoid and nonlymphoid organs, Science 350 (2015) 981–985. 10.1126/science.aac9593.

[2] K.R. Bartemes, K. Iijima, T. Kobayashi, G.M. Kephart, A.N. McKenzie, H. Kita, IL-33– Responsive Lineage−CD25+CD44hi Lymphoid Cells Mediate Innate Type 2 Immunity and Allergic Inflammation in the Lungs, J Immunol 188 (2012) 1503–1513. 10.4049/jimmunol.1102832.

[3] L.A. Monticelli, L.C. Osborne, M. Noti, S.V. Tran, D.M.W. Zaiss, D. Artis, IL-33 promotes an innate immune pathway of intestinal tissue protection dependent on amphiregulin–EGFR interactions, Proceedings of the National Academy of Sciences 112 (2015) 10762–10767. 10.1073/pnas.1509070112.

[4] D.R. Neill, S.H. Wong, A. Bellosi, R.J. Flynn, M. Daly, T.K.A. Langford, C. Bucks, C.M. Kane, P.G. Fallon, R. Pannell, H.E. Jolin, A.N.J. McKenzie, Nuocytes represent a new innate effector leukocyte that mediates type-2 immunity, Nature 464 (2010) 1367– 1370. 10.1038/nature08900.

[5] R.E. Mebius, T. Miyamoto, J. Christensen, J. Domen, T. Cupedo, I.L. Weissman, K. Akashi, The Fetal Liver Counterpart of Adult Common Lymphoid Progenitors Gives Rise to All Lymphoid Lineages, CD45+CD4+CD3− Cells, As Well As Macrophages, The Journal of Immunology 166 (2001) 6593–6601. 10.4049/jimmunol.166.11.6593.

[6] C. Possot, S. Schmutz, S. Chea, L. Boucontet, A. Louise, A. Cumano, R. Golub, Notch signaling is necessary for adult, but not fetal, development of RORγt+ innate lymphoid cells, Nat Immunol 12 (2011) 949–958. 10.1038/ni.2105.

[7] S.H. Wong, J.A. Walker, H.E. Jolin, L.F. Drynan, E. Hams, A. Camelo, J.L. Barlow, D.R. Neill, V. Panova, U. Koch, F. Radtke, C.S. Hardman, Y.Y. Hwang, P.G. Fallon, A.N.J. McKenzie, Transcription factor RORα is critical for nuocyte development, Nat Immunol 13 (2012) 229–236. 10.1038/ni.2208.

[8] X. Yu, Y. Wang, M. Deng, Y. Li, K.A. Ruhn, C.C. Zhang, L.V. Hooper, The basic leucine zipper transcription factor NFIL3 directs the development of a common innate lymphoid cell precursor, eLife 3 (2014) e04406. 10.7554/eLife.04406.

[9] Q. Yang, F. Li, C. Harly, S. Xing, L. Ye, X. Xia, H. Wang, X. Wang, S. Yu, X. Zhou, M. Cam, H.-H. Xue, A. Bhandoola, TCF-1 upregulation identifies early innate lymphoid progenitors in the bone marrow, Nat Immunol 16 (2015) 1044–1050. 10.1038/ni.3248.

[10] C.S.N. Klose, M. Flach, L. Möhle, L. Rogell, T. Hoyler, K. Ebert, C. Fabiunke, D. Pfeifer, V. Sexl, D. Fonseca-Pereira, R.G. Domingues, H. Veiga-Fernandes, S.J. Arnold, M. Busslinger, I.R. Dunay, Y. Tanriver, A. Diefenbach, Differentiation of Type 1 ILCs from a Common Progenitor to All Helper-like Innate Lymphoid Cell Lineages, Cell 157 (2014) 340–356. 10.1016/j.cell.2014.03.030.

[11] C. Harly, M. Cam, J. Kaye, A. Bhandoola, Development and differentiation of early innate lymphoid progenitors, Journal of Experimental Medicine 215 (2018) 249–262. 10.1084/jem.20170832.

[12] M.G. Constantinides, B.D. McDonald, P.A. Verhoef, A. Bendelac, A committed precursor to innate lymphoid cells, Nature 508 (2014) 397–401. 10.1038/nature13047.

[13] T. Hoyler, C.S.N. Klose, A. Souabni, A. Turqueti-Neves, D. Pfeifer, E.L. Rawlins, D. Voehringer, M. Busslinger, A. Diefenbach, The transcription factor GATA-3 controls cell fate and maintenance of type 2 innate lymphoid cells, Immunity 37 (2012) 634–648. 10.1016/j.immuni.2012.06.020.

[14] K. Moro, T. Yamada, M. Tanabe, T. Takeuchi, T. Ikawa, H. Kawamoto, J.-I. Furusawa, M. Ohtani, H. Fujii, S. Koyasu, Innate production of T(H)2 cytokines by adipose tissue-associated c-Kit(+)Sca-1(+) lymphoid cells, Nature 463 (2010) 540–544. 10.1038/nature08636.

[15] N. Satoh-Takayama, S. Lesjean-Pottier, P. Vieira, S. Sawa, G. Eberl, C.A.J. Vosshenrich, J.P. Di Santo, IL-7 and IL-15 independently program the differentiation of intestinal CD3−NKp46+ cell subsets from Id2-dependent precursors, J Exp Med 207 (2010) 273–280. 10.1084/jem.20092029.

[16] Y. Yokota, A. Mansouri, S. Mori, S. Sugawara, S. Adachi, S. Nishikawa, P. Gruss, Development of peripheral lymphoid organs and natural killer cells depends on the helix-loop-helix inhibitor Id2, Nature 397 (1999) 702–706. 10.1038/17812.

[17] M.D. Boos, Y. Yokota, G. Eberl, B.L. Kee, Mature natural killer cell and lymphoid tissue–inducing cell development requires Id2-mediated suppression of E protein activity, The Journal of Experimental Medicine 204 (2007) 1119–1130. 10.1084/jem.20061959.

[18] M. Ji, H. Li, H.C. Suh, K.D. Klarmann, Y. Yokota, J.R. Keller, Id2 intrinsically regulates lymphoid and erythroid development via interaction with different target proteins, Blood 112 (2008) 1068–1077. 10.1182/blood-2008-01-133504.

[19] J.A. Walker, P.A. Clark, A. Crisp, J.L. Barlow, A. Szeto, A.C.F. Ferreira, B.M.J. Rana, H.E. Jolin, N. Rodriguez-Rodriguez, M. Sivasubramaniam, R. Pannell, J. Cruickshank, M. Daly, L. Haim-Vilmovsky, S.A. Teichmann, A.N.J. McKenzie, Polychromic Reporter Mice Reveal Unappreciated Innate Lymphoid Cell Progenitor Heterogeneity and Elusive ILC3 Progenitors in Bone Marrow, Immunity 51 (2019) 104–118.e7. 10.1016/j.immuni.2019.05.002.

[20] W. Xu, D.E. Cherrier, S. Chea, C. Vosshenrich, N. Serafini, M. Petit, P. Liu, R. Golub, J.P. Di Santo, An Id2RFP-reporter mouse redefines innate lymphoid cell precursor potentials, Immunity 50 (2019) 1054–1068.

[21] M.A. Cannarile, N.A. Lind, R. Rivera, A.D. Sheridan, K.A. Camfield, B.B. Wu, K.P. Cheung, Z. Ding, A.W. Goldrath, Transcriptional regulator Id2 mediates CD8+ T cell immunity, Nat Immunol 7 (2006) 1317–1325. 10.1038/ni1403.

[22] Y.-Y. Lin, M.E. Jones-Mason, M. Inoue, A. Lasorella, A. Iavarone, Q.-J. Li, M.L. Shinohara, Y. Zhuang, Transcriptional regulator Id2 is required for the CD4 T cell immune response in the development of experimental autoimmune encephalomyelitis, J Immunol 189 (2012) 1400–1405. 10.4049/jimmunol.1200491.

[23] H. Sun, J. Gao, P. Zhang, Y. Wang, Y. Chang, J. Miao, P. Zhu, [Increased expression of inhibitor of differentiation 2 (Id2) in CD4+T cells in joint synovial fluid of patients with rheumatoid arthritis], Xi Bao Yu Fen Zi Mian Yi Xue Za Zhi 38 (2022) 333–337.

[24] Q.P. Nguyen, K.K. Takehara, T.Z. Deng, S. O’Shea, M. Heeg, K.D. Omilusik, J.J. Milner, S. Quon, M.E. Pipkin, J. Choi, S. Crotty, A.W. Goldrath, Transcriptional programming of CD4+ TRM differentiation in viral infection balances effector⁻ and memory-associated gene expression, Sci Immunol 8 (2023) eabq7486. 10.1126/sciimmunol.abq7486.

[25] X. Wang, J. Li, L. Rebuffet, M. Cheng, B. Bao, Y. Chen, X. Zheng, Y. Chen, H. Sun, R. Sun, E. Vivier, H. Peng, Z. Tian, Innate lymphoid cells originate from fetal liver–derived tissue-resident progenitors, Science Immunology 10 (2025) eadu7962. 10.1126/sciimmunol.adu7962.

[26] P. Naef, C.A. Jaeger-Ruckstuhl, N. Schnüriger, S. Forster, I. Monteiro, D. Brigger, A. Eggel, K. Kessenbrock, C. Riether, A.F. Ochsenbein, IL-33/ST2 signaling in ILC2s drives exhaustion and myeloid skewing of HSCs in response to hematopoietic stress and aging, iScience 28 (2025) 112378. 10.1016/j.isci.2025.112378.

[27] T. Sudo, Y. Motomura, D. Okuzaki, T. Hasegawa, T. Yokota, J. Kikuta, T. Ao, H. Mizuno, T. Matsui, D. Motooka, Group 2 innate lymphoid cells support hematopoietic recovery under stress conditions, Journal of Experimental Medicine 218 (2021).

[28] S. Chea, C. Possot, T. Perchet, M. Petit, A. Cumano, R. Golub, CXCR6 Expression Is Important for Retention and Circulation of ILC Precursors, Mediators of Inflammation 2015 (2015) 1–15. 10.1155/2015/368427.

[29] S. Chea, S. Schmutz, C. Berthault, T. Perchet, M. Petit, O. Burlen-Defranoux, A.W. Goldrath, H.-R. Rodewald, A. Cumano, R. Golub, Single-cell gene expression analyses reveal heterogeneous responsiveness of fetal innate lymphoid progenitors to notch signaling, Cell Reports 14 (2016) 1500–1516.

[30] T. Kawamoto, M. Shimizu, A method for preparing 2⁻ to 50-µm-thick fresh-frozen sections of large samples and undecalcified hard tissues, Histochem Cell Biol 113 (2000) 331–339. 10.1007/s004180000149.

[31] E. Gabandé-Rodríguez, G. Soto-Heredero, E. Carrasco, C. Anerillas, J.I. Escrig-Larena, J. Martínez-Cano, S. Delgado-Pulido, I. Francos-Quijorna, G. Desdín-Micó, T. Smirnova, M.M. Gómez De Las Heras, Á. Fernández-Almeida, A. Leandri, A. Cugurra, E.M. Blanco, E. Winand-Osete, P. Ramírez-Ruiz De Erenchun, V. Zorita, C. Bean, V. Basset, F. Solly, A. Garrido, R. De Cabo, S. Mañes, M. Pasparakis, M. Iannacone, C. Cobaleda, O. Naveiras, M. Gorospe, M. Mittelbrunn, Cytotoxic CD4+ T cells induce age-associated myelopoiesis through CCL5–CCR5 signaling, Nat Aging 6 (2026) 2110– 2125. 10.1038/s43587-026-01209-9.

[32] H. Maazi, N. Patel, I. Sankaranarayanan, Y. Suzuki, D. Rigas, P. Soroosh, G.J. Freeman, A.H. Sharpe, O. Akbari, ICOS:ICOS-Ligand Interaction Is Required for Type 2 Innate Lymphoid Cell Function, Homeostasis, and Induction of Airway Hyperreactivity, Immunity 42 (2015) 538–551. 10.1016/j.immuni.2015.02.007.

[33] D. Paclik, C. Stehle, A. Lahmann, A. Hutloff, C. Romagnani, ICOS regulates the pool of group 2 innate lymphoid cells under homeostatic and inflammatory conditions in mice, Eur J Immunol 45 (2015) 2766–2772. 10.1002/eji.201545635.

